# Thiosulfate-hydrogen peroxide redox oscillator as pH driver for ribozyme activity in the RNA world

**DOI:** 10.1101/020693

**Authors:** Rowena Ball, John Brindley

## Abstract

The RNA world of more than 3.7 billion years ago may have drawn on thermal and pH oscillations set up by the oxidation of thiosulfate by hydrogen peroxide (the THP oscillator) as a power source to drive replication. Since this primordial RNA also must have developed enzyme functionalities, in this work we examine the responses of two simple ribozymes to a THP periodic drive, using experimental rate and thermochemical data in a dynamical model for the coupled, self-consistent evolution of all reactants and intermediates. The resulting time traces show that ribozyme performance can be enhanced under pH cycling, and that thermal cycling may have been necessary to achieve large performance gains. We discuss three important ways in which the dynamic hydrogen peroxide medium may have acted as an agent for development of the RNA world towards a cellular world: proton gradients, resolution of the ribozyme versus replication paradox, and vesicle formation.

## Introduction

One of the major open questions pertaining to the origin and evolution of life concerns the power source for the RNA world (Higgs and Lehman 2015). Of course, the primary power source presumably was solar or geothermal, but those sources acting directly on the pre-cellular RNA world cannot provide the periodic drive that is vital for sustaining and evolving a non-cellular living system. It is now accepted that the RNA world must have been subject to a thermal drive with a period of order 100 s in order to survive, replicate and amplify (Szostak 2012; Ball and Brindley 2014; Kreysing et al 2015), but cellular life is powered by proton gradients — the ubiquitous proton motive force (Mitchell 1961; Lane et al 2010). How might thermal cycling have given way to proton gradients?

We have proposed that hydrogen peroxide may have acted as a power source and agent for change in the pre-cellular RNA world. As the thiosulfate-hydrogen peroxide (THP) thermochemical oscillator it can provide a periodic thermal and pH drive for RNA replication (Ball and Brindley 2014, 2015a), and its physical properties and structure, especially its axial chirality, may have helped to mediate homochiral proto-life (Ball and Brindley 2015b). In this work we report the results of fully self-consistent simulations in which the THP oscillator is used to drive RNA enzyme, or ribozyme, reactions, where the rate constants for the RNA reactions are both temperature and pH dependent. We find that pH cycling improves the performance of the ribozyme reactions, and can resolve the ‘replication versus ribozyme activity’ paradox (Ivica et al 2013). We discuss the provision of a proton motive force by the THP oscillator, in a spatially extended, bounded system, and exemplify the potential for the THP oscillator to perfom multiple, parallel functions that may have facilitated the development of cell-based life that could prosper as the supply of hydrogen peroxide diminished.

Hammerhead ribozymes are a much-studied class of small RNA motifs that undergo autocatalytic self-cleavage (Scott et al 2013). They were discovered first in pathogenic plant viroids, but hammerhead complementary sequences have now been detected in the genomes of most phyla, and they are believed to be molecular survivors from the RNA world (de la Peña and García-Robles 2010). For this latter reason we consider that their performance under a THP oscillatory drive is a question of some importance.

The THP oscillator has been studied for many years by two camps of researchers, which apparently remained entirely separate. The first group was based in chemical engineering, and focussed exclusively on the simple thermal oscillations produced by oxidation of thiosulfate by hydrogen peroxide at high concentrations (e.g., Chang and Schmitz (1975); Grau et al (2000)). The second group was based in organic chemistry, and focussed exclusively on the (more or less) complex pH oscillations under isothermal conditions produced by this reaction system at low concentrations (e.g. Orbán and Epstein (1987); Rábai and Hanazaki (1999a)). Although high quality experimental data were obtained by both camps, some of which we make use of in this work, we believe that this narrow, *intra*-disciplinary development of the field meant that the versatility and potential of the THP reaction system was under-appreciated. In none of the works by either group was the suggestion made that the THP oscillator could drive natural or technological processes. Thus one of the general achievements of our work is unification of these two strands.

Other pH oscillators have been used experimentally to drive biological processes, so there is every reason to suppose the THP oscillator may have driven ribozyme activity in the RNA world. For example, Lagzi et al (2010) used the methylene glycol-sulfite-gluconolactone (MGSG) oscillator to interconvert vesicles and micelles, Liedl and Simmel (2005) and Liedl et al (2006) used the oscillatory oxidation of sulfite and thiosulfate by iodate to drive DNA conformation changes, and Qi et al (2013) used the iodate-thiosulfate-ferrocyanide oscillatory pH system to drive switchable DNA machines.

In the following sections we describe the THP reaction system and the dynamical model, present and discuss results for reactions of two pH-dependent hammerhead ribozymes, and discuss some other possible effects of the THP oscillator on the RNA world.

## Model and methods

**Table 1.**
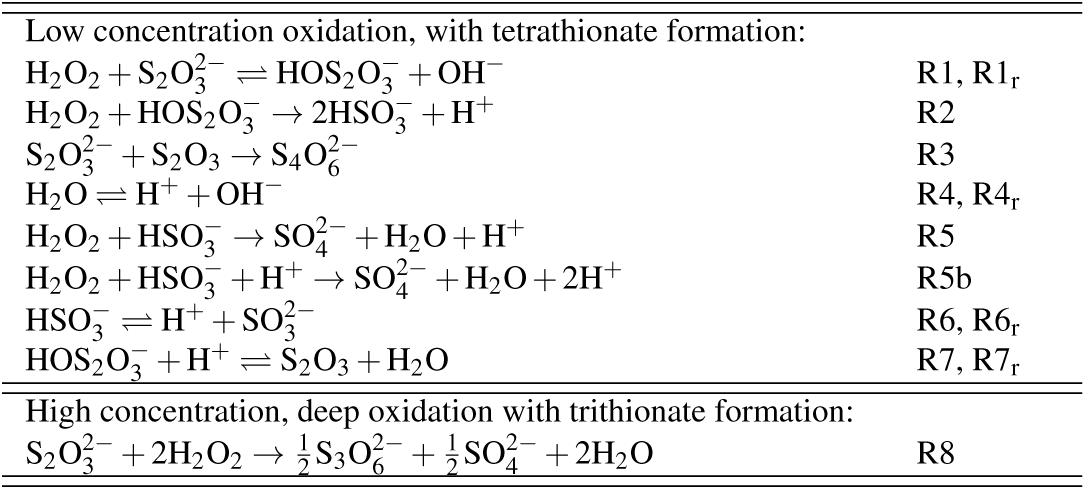
Reactions involved in the THP thermo-pH oscillator. Low concentration oxidation, with tetrathionate formation:

|  |  |  |
| --- | --- | --- |
| Low concentration oxidation, with tetrathionate formation: |  |  |
| $\text{H}_2\text{O}_2 + \text{S}_2\text{O}_3^{2-} \rightleftharpoons \text{HOS}_2\text{O}_3^- + \text{OH}^-$ | | R1, R1<sub>r</sub> |
| $\text{H}_2\text{O}_2 + \text{HOS}_2\text{O}_3^- \rightarrow 2\text{HSO}_3^- + \text{H}^+$ | | R2 |
| $\text{S}_2\text{O}_3^{2-} + \text{S}_2\text{O}_3 \rightarrow \text{S}_4\text{O}_6^{2-}$ | | R3 |
| $\text{H}_2\text{O} \rightleftharpoons \text{H}^+ + \text{OH}^-$ | | R4, R4<sub>r</sub> |
| $\text{H}_2\text{O}_2 + \text{HSO}_3^- \rightarrow \text{SO}_4^{2-} + \text{H}_2\text{O} + \text{H}^+$ | | R5 |
| $\text{H}_2\text{O}_2 + \text{HSO}_3^- + \text{H}^+ \rightarrow \text{SO}_4^{2-} + \text{H}_2\text{O} + 2\text{H}^+$ | | R5b |
| $\text{HSO}_3^- \rightleftharpoons \text{H}^+ + \text{SO}_3^{2-}$ | | R6, R6<sub>r</sub> |
| $\text{HOS}_2\text{O}_3^- + \text{H}^+ \rightleftharpoons \text{S}_2\text{O}_3 + \text{H}_2\text{O}$ | | R7, R7<sub>r</sub> |
| High concentration, deep oxidation with trithionate formation: |  |  |
| $\text{S}_2\text{O}_3^{2-} + 2\text{H}_2\text{O}_2 \rightarrow \frac{1}{2}\text{S}_3\text{O}_6^{2-} + \frac{1}{2}\text{SO}_4^{2-} + 2\text{H}_2\text{O}$ | | R8 |

Reactions and equilibria R1 to R7_r_ in Table 1 describe the mechanism that was proposed by Rábai and Hanazaki (1999a) for the THP pH oscillator. This mechanism operates at low feed concentrations (*∼*mM), for which the major sulfur species product is tetrathionate. Under these conditions the reaction system remains nearly isothermal, because the specific reaction heat is small and can be absorbed by quantized intra- and intermolecular modes, or dissipated.

At high feed concentrations (*∼*M) reaction R8 in Table 1 appears to take over (Chang and Schmitz 1975; Grau et al 2000), with production of the trithionate and suppression of tetrathionate production. In this regime the THP oscillator is described by reaction R8 and the acid-base equilibria R4, R4_r_ and R6, R6_r_, and both pH and thermal oscillations of substantial amplitude can occur. The thermal and pH oscillations in both regimes, and a smooth transition from one to the other, were characterised in Ball and Brindley (2014, 2015a).

Motivated by the likely occurrence on the early Earth of situations in which ‘primordial soup’ flowed through porous rocks, we use a well stirred continuous flow (CSTR) paradigm, in which the RNA species are carried as a dilute suspension in an ambient steady flow of chemically reactive fluid, which can support thermochemical oscillations (Ball and Brindley 2014). Each pore constitutes a cell in thermal contact with its environment, and we assume that individual pores in some macroscopic sized matrix are connected by narrow channels which permit ready throughput of the ambient fluid (and small monomeric species) but which offer hindrance to the passage of polymers, increasingly so with increasing strand length. The transport of flexible coiled or otherwise configured polymers through narrow channels is of course a highly complex process (Deen 1987), much studied as a key part of many industrial processes, especially in the pharmaceutical field, and medical applications. Here we content ourselves with assuming simply that there will be trapping and accumulation in pores of the ribozymes and other large molecules. The thermoconvective experiments and simulations of Braun’s group (e.g., Baaske et al (2008); Kreysing et al (2015) have demonstrated preferential accumulation and retention of long nucleotide chains in open pore systems. Such accumulations are favorable for replication (treated in the companion paper, Ball and Brindley (2015a)) and ribozyme activity, enhanced by the macroscopic scale THP oscillations which, under appropriate conditions, exhibit themselves as travelling waves through the porous matrix. At the scale of the individual (microscopic) pore, the passage of each wave is of course seen as a purely temporal oscillation.

Our dynamical model is based on the rate laws for the reactions in Table 1, as well as the rate laws for ribozyme reactions described in the next section, including the temperature dependence of the rate coefficients. The enthalpy balance includes all reaction enthalpy contributions, the heat transported by the flow, and the rate of heat lost to the environment:

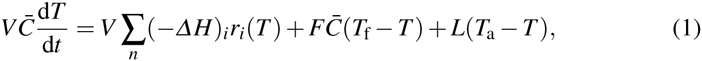

where the summation is over *n* reaction rates *r*_*i*_(*T*). A mass balance for each reactant species has the form

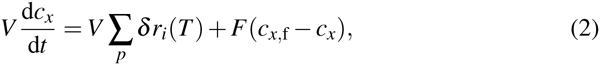

where *c*_*x*_ is the concentration of reactant or intermediate *x*, the summation is over the *p* reaction rates that involve *x*, *c*_*x,f*_ = 0 for intermediate species, *δ* = 1 for intermediates that are produced and *δ* = −1 for reactants and intermediates that are consumed. The rate coefficients *k*_*i*_ of the reaction rates *r*_*i*_ have Arrhenius temperature dependence:

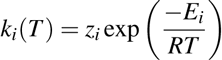

The rate coefficients for ribozyme reactions depend on pH, as well as temperature, as given in the next section. The symbols and quantities used here and in the rest of the paper are defined in table 8, where numerical values for the non-variable parameters are also given.

**Table 2.**
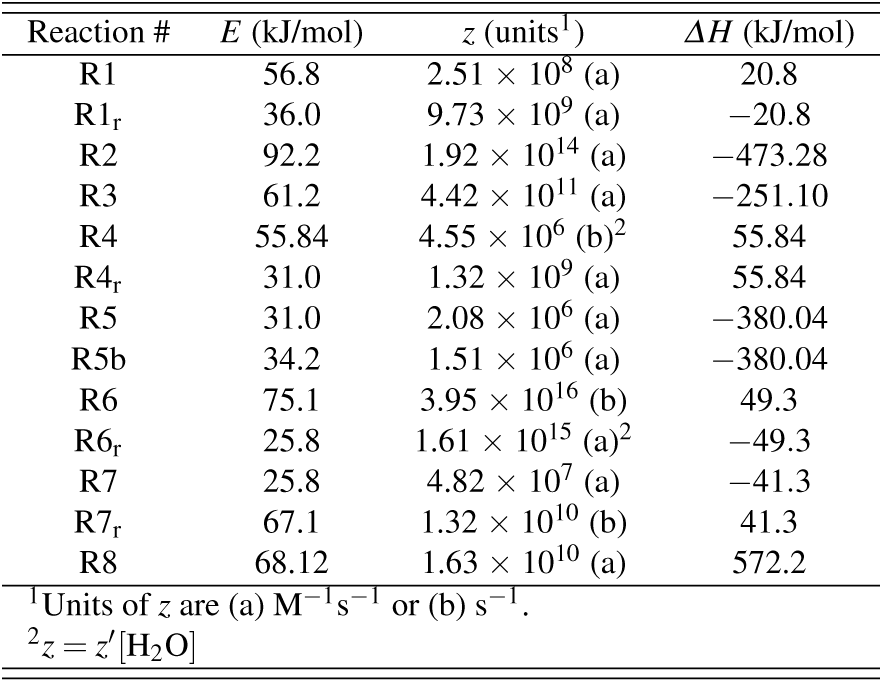
Rate parameters and reaction enthalpies for the THP reactions in Table 1 (Goldberg and Parker 1985; Williamson and Rimstidt 1992; Rábai and Hanazaki 1999a).

The CSTR equations (1) and (2) may be thought of as an idealized model of the proposed physical setting for the RNA-THP oscillator system. The volume *V* represents that of a rock pore in a submarine hydrothermal system, as described above. Highly porous mineral precipitates around hydrothermal vents have been widely accepted as providing the necessary open thermodynamic, far from equilibrium, conditions, as well as surface and mineral catalysts and components such as reduced sulfoxyions (Baaske et al 2008; Lane and Martin 2012; Kreysing et al 2015). The prebiotic sources of hydrogen peroxide were reviewed in Ball and Brindley (2014).

Using the kinetic data and reaction enthalpies in Tables 2, 3 and 5, and selected values for the feed concentrations *c*_*x,f*_ and flow rate *F*, we integrated Eqs (1) and (2) from convenient initial conditions using a stiff integrator to obtain time series. We also post-processed oscillatory time series to obtain accurate averaged quantities as follows: The time trajectory for each relevant quantity was integrated numerically over at least 20 cycles using the trapezoidal rule on 20,000 data points for high accuracy, then divided by the total number of seconds to obtain true weighted average values of the oscillating quantities. These weighted averages are denoted as 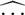 hatted quantities. (Simply taking arithmetic time averages yields incorrect and misleading values, especially as the oscillations become less sinusoidal.)

**Table 3.**
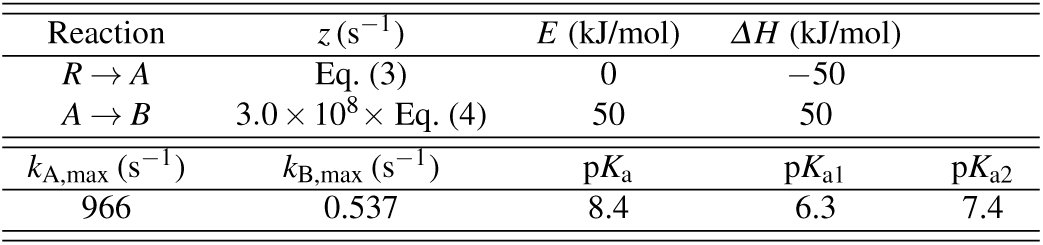
Kinetic parameters, reaction enthalpies, and dissociation constants for R9.

| Reaction | $z$ (s<sup>-1</sup>) | $E$ (kJ/mol) | $\Delta H$ (kJ/mol) | | |
| --- | --- | --- | --- | --- | --- |
| $R \rightarrow A$ | Eq. (3) | 0 | -50 | | |
| $A \rightarrow B$ | $3.0 \times 10^8 \times \text{Eq. (4)}$ | 50 | 50 | | |
| $k_{A,\max}$ (s<sup>-1</sup>) | $k_{B,\max}$ (s<sup>-1</sup>) | $pK_a$ | $pK_{a1}$ | $pK_{a2}$ | |
| 966 | 0.537 | 8.4 | 6.3 | 7.4 |  |

## Results and discussion

Example I: Ribozyme activity in the low feed concentration pH oscillator regime

Buskiewicz and Burke (2012) studied experimentally the pH-dependent kinetics of folding and cleavage of variants of a 35 bp hammerhead ribozyme. The reaction can be regarded as proceeding in two sequential steps:

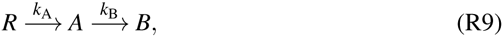

where *R* is the inactive, unfolded ribozyme, *A* is the correctly folded, active ribozyme, and *B* is the self-cleavage product. The experimental rate constant for folding was fitted to the following pH dependence:

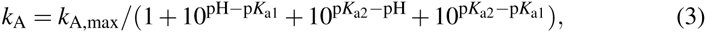

and that for self-cleavage was found to depend on only one titratable group:

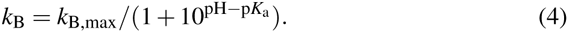

In these dependences *k*_A, max_, *k*_B, max_ are the the rate constants at maximum pH, and *K*_a_, *K*_a1_, *K*_a2_ are acid dissociation constants of titratable groups on the ribozyme.

We used the THP oscillator in the low feed concentration, pH regime to drive reactions R9, using the above pH dependences for the unmodified ribozyme (G8C3) in Buskiewicz and Burke (2012). Accordingly, Eqs (1) and (2) were set up using the rates and reactant/intermediate concentrations for reactions R1, R1_f_ to R7, R7_f_ in Table 1 and reactions R9. Values of the thermokinetic parameters and reaction enthalpies for reactions R9, which we estimated, and the isothermal kinetic parameters from Buskiewicz and Burke (2012), are given in Table 3.

A time series resulting from the computations is shown in Fig. 1, rendered in terms of the pH (a.), concentration of folded species *A* (b.), concentration of cleaved product *B* (c.), and the temperature (d.). In e. we have superimposed the time traces from a., b. c. and d. over one period, to facilitate interpretation of the phase relationships. The following points help to interpret Fig. 1:

**–** The oscillations are complex in form — in general this is expected, since we are working with a high-dimensional dynamical system. Complex dynamics, such as chaotic and quasiperiodic behaviour, have been recorded experimentally in the THP system (Rábai and Hanazaki 1999b; Bakesš et al 2008).
**–** The pH amplitude varies between a minimum of 4.3 and a maximum of 7.5 and the integrated weighted average pH is 5.19. From Buskiewicz and Burke (2012) the rate constant for folding, *k*_A_, is highest at pH values between 6.5 and 7.5, a region in Fig. 1a. which is sampled in one burst of about 60 s in each period of 250 s.
**–** We note in b. that the folded species *A* is favoured at higher pH and in c. the self-cleaved product *B* is favoured at lower pH.
**–** The temperature oscillations d. would be undetectable in an experiment; nevertheless it is interesting to note in e. that the temperature rises abruptly as the pH falls — this is due largely to production of hydrogen ions by reaction R2 in Table 1, which is highly exothermic, Table 2.

**Fig. 1.**
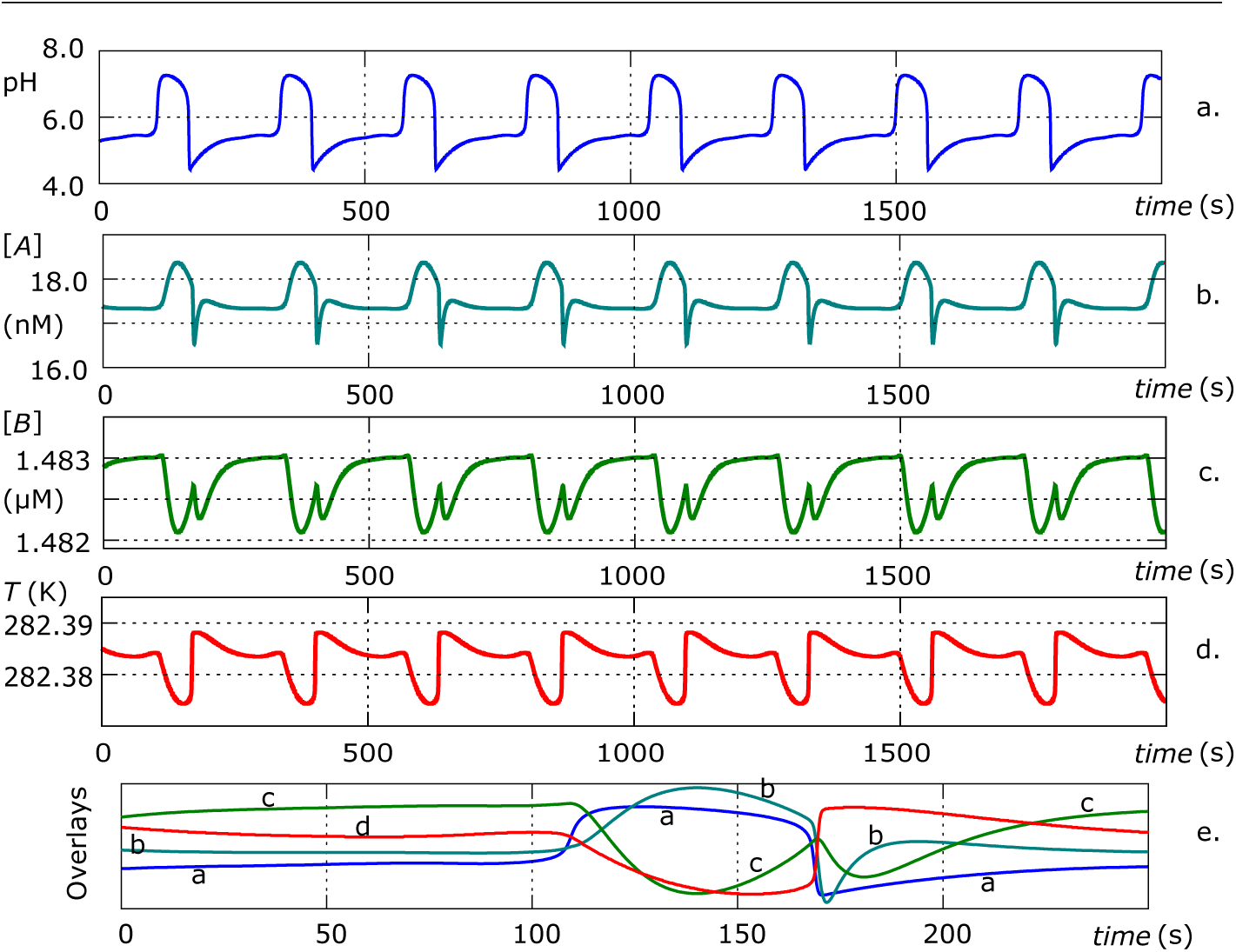
Time series for the THP system in the pH regime with reactions R9 included self-consistently. *F* = 0.01061 *μ*l s^-1^, [*R*]_f_ = 1.5 *μ*M, [H_2_O_2_]_f_ = 0.0135 M, 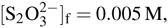, [H^+^]_f_ = 0.5 mM, 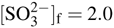, *L* = 0.1 mW, *t* = 250 s. Traces a., b., c. and d. over one period are overlaid in e.

Does the system under oscillatory drive exhibit better performance than the system under constant pH? Unavoidably, we can only define ‘better performance’ in anthropocentric terms, so for illustrative purposes we define a better performing system as having lower concentration of *A* in the reaction cell, because we want more of it to self-cleave to product *B*, and higher concentration of *B*, since that would indicate higher ribozyme productivity. Whether this narrow definition represents better performance in the context of the RNA world is unknown, but, as we shall see, the results can provide an indicative answer to the more general question: Is the RNA world *fitter* under an oscillatory pH drive?

**Table 4.**
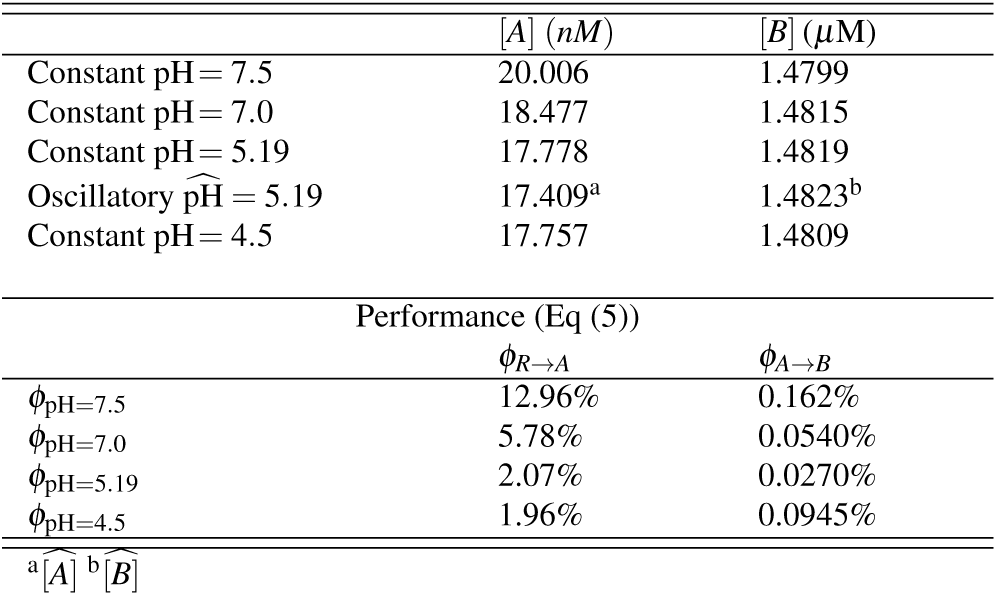
Comparison of the performance of reactions R9 under THP oscillatory and constant pH drive.

**Fig. 2.**
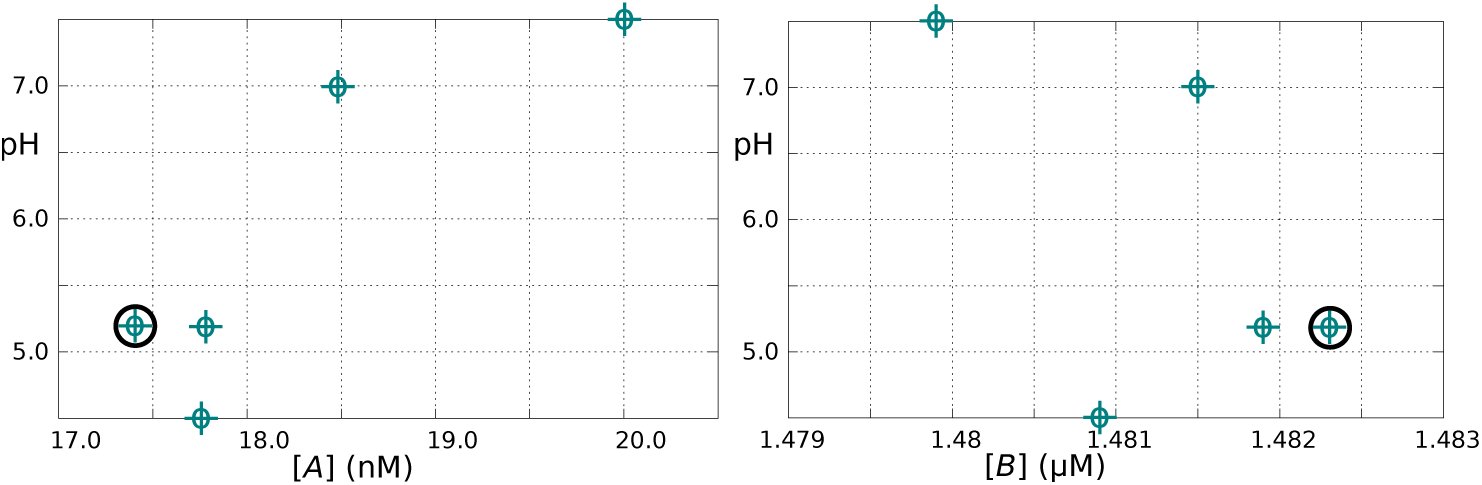
Plots of data points in Table 4. The data points obtained from the oscillatory time series as integrated instantaneous quantities are circled.

To compare the performance of the system under oscillatory and constant pH drive we carried out some runs at constant pH (i.e., using only reactions R4, R4_r_, R6 and R6_r_ from Table 1) at the parameter values given in the caption to Fig. 1 and recorded the steady-state concentrations, and made use of the integrated weighted average quantities, as described earlier, from the oscillatory time series plotted in Fig. 1. As measures of performance we define

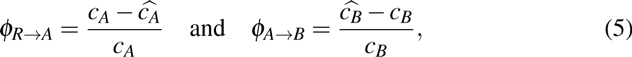

where *c*_*A*_and *c*_*B*_are steady state concentrations under constant pH and 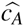 and 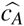 are integrated weighted average concentrations under oscillatory drive. Results are given in Table 4 and plotted in Fig. 2.

Conversion to cleaved product *B* is generally high, since the feed concentration of ribozyme *R* is 1.5 *μ*M. There is a trend towards lower concentration of folded species *A* and higher concentration of cleaved product *B* as the pH is decreased. The exception is the concentration of *B* at pH 4.5, which has decreased sharply. The hatted data points are highlighted by circles. In each case the effect of the oscillatory pH drive is to exaggerate the trend. Considering the performance results in Table 4, the oscillatory pH drive achieves a moderate increase in *ϕ_R→A_* which decreases with pH, and a small increase in *ϕ_A→B_* which decreases with pH, except for pH 4.5, for which it increases. (We note that results for pH *<* 5 may not be reliable, since pH 5 was the limit of experimental measurements (Buskiewicz and Burke 2012).)

Can performance gains of only 0.027–0.162% be considered significant in terms of the RNA world? We would argue that changes of all kinds in the RNA world must be small, because large perturbations are more likely to wipe out life, or incipient life, than mediate and enhance it. A large, rapid gain in the performance of a ribozyme must come at a price, at the expense of a (possibly catastrophic) loss of some other capability of the RNA world, such as replicative capacity. This is simply an extension of a general principle of evolutionary biology, that gains in fitness are made by small, incremental changes.

Small differences in concentration can lead to large differences in abundance. If the ribozyme can template and self-replicate, the dynamics will ensure that the fittest species will rapidly dominate in large excess. In the current context the ‘fittest species’ is that which thrives or performs better under pH cycling than other species.

Further, the definition of performance, Eq. (5), and the results in Figs 1 and 2 and Table 4 need not be taken too literally or narrowly. What they *do* suggest is that an oscillatory pH drive allows the system to access many more degrees of freedom and gives it the flexibility potentially to adapt, self-optimize, and run multiple operations simultaneously. For example, the ribozyme may develop the ability to carry out at several functions, each of which operates optimally at a different value of the pH in a complex cycle. In the RNA world the ribozyme must also undergo replication, a process for which the optimum rate may occur at a lower pH than that for enzyme functionality. The work of Cafferty et al (2014), using monomers that mimic nucleic acid base pairing, has shown that supramolecular polymers can exhibit sharp transitions to different behaviours under pH control, and it is noted that this might provide clues as to how the first genetic polymers responded to periodic changes in pH. Under constant pH the ribozyme cannot exhibit such versatility. In this sense the RNA world under oscillatory drive resembles a truly living system.

Since the assigned activation energy for this reaction, 50 kJ/mole, is not insignificant we suspect that this system may perform better in the high feed concentration, thermo-pH regime, which was characterised in Ball and Brindley (2015a), and in the following section we test this idea using data for another ribozyme.

**Table 5.** Kinetic and thermochemical parameters derived from data in Hertel and Uhlenbeck (1995).

|  | cleavage | ligation |
| --- | --- | --- |
| $z_0$ ($\text{s}^{-1}$) | $9.8 \times 10^{13*}$ | $5.0 \times 10^{7*}$ |
| $E$ (kJ/mol) | 92 | 42 |
| $m$ | 1.2 | 1.0 |
| $\Delta H$ (kJ/mol) | 50 | -50 |

\*assigned values

Example II: Ribozyme cleavage-ligation equilibrium

A hammerhead ribozyme studied by Hertel and Uhlenbeck (1995) was found to undergo pH-dependent self-cleavage and self-ligation reactions, which may be represented by the following equilibrium:

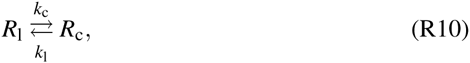

where *R*_l_ and *R*_c_ represent the ligated and cleaved ribozyme respectively, and the equilibrium strongly favours cleavage over ligation. Therefore it is a good candidate for evaluation under THP oscillatory drive. Hertel and Uhlenbeck (1995) found a power-law dependence of the rate constants for reactions R10 on the hydrogen ion concentration, and from their data the rate constants may be expressed as

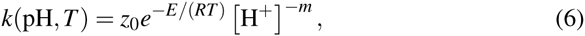

Numerical values for the rate parameters and reaction enthalpies are given in Table 5.

First, we used the THP oscillator in the low feed concentration, pH oscillatory regime to drive reactions R10 under nearly isothermal conditions, so Eqs (1) and (2) were set up using the rates and reactant/intermediate concentrations for reactions R1, R1_f_ to R7, R7_f_ in Table 1 and reactions R10. The computed time series is shown in Fig. 3, rendered in terms of the pH and the concentrations of ligated ribozyme [*R*_l_] and cleaved product [*R*_c_].

We find that the yield of ligated product is indeed low and that of cleaved product is high, as was also found by Hertel and Uhlenbeck (1995); from the computed data file, we derive the integrated average yield of ligated product as 1.14%. What is the effect of the oscillatory pH drive on yields, compared with a constant pH drive? This comparison is made in Table 6. The values for integrated weighted average quantities 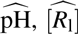 and 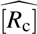 for two runs are listed, and the concentrations are compared with those obtained by running the system at the constant pH corresponding. Evidently the effect of pH cycling is to drive the equilibrium further towards the cleaved product, with 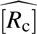 2.3% higher at 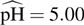 and 7.0% higher at 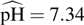. Whether this effect is ‘good’ or ‘bad’ depends entirely on which product — cleavage or ligation — benefits the RNA world more. However, since the reactant perforce is a modern ribozyme, optimised by evolution for efficient cleavage, we can say with confidence that pH cycling enhances its action.

On the other hand, if it is desired to improve ligation yield it may be advantageous to run the reactions R10 in the high concentration, thermo-pH oscillatory regime of the THP oscillator. In this case production of the tetrathionate is suppressed and the dynamical system, set up with Eqs (1) and (2), uses the rates and reactant/intermediate concentrations for reaction R8, and the pH equilibria R4, R4_r_ and R6, R6_r_ in Table 1 and reactions R10. The results are presented in Fig. 4 and Table 7. The yield of ligation product is 13.4%, improved by a factor of 4 over that for 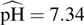 in Table 6. What is the effect of thermal cycling on yields, compared to driving the system at a constant temperature of 298 K, as used by Hertel and Uhlenbeck (1995)? This, too, is shown in Table 7. The constant pH in the thermostatted system is adjusted to 7.55 and the production of cleaved product is favoured by 4.1% over that for the thermal cycling system. From another point of view we can say that thermal cycling improves the yield of ligated product by 36%.

**Fig. 3.**
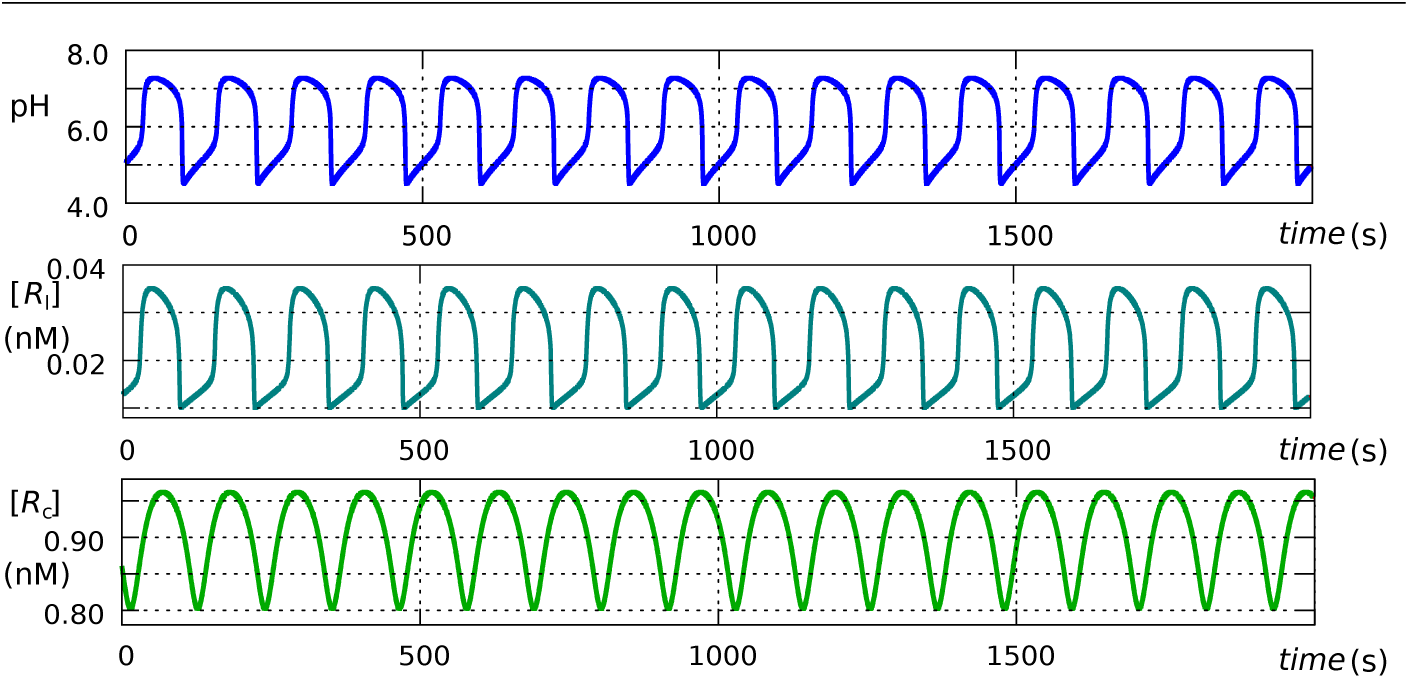
The yield of ligated product is low under nearly-isothermal oscillatory pH drive. *F* = 0.0090 *μ*l s^-1^, [*R*_c_]_f_ = 1.1 nM, [H_2_O_2_]_f_ = 0.0135 M, 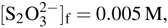, [H^+^]_f_ = 0.5 mM, 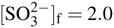, period=125 s.

**Table 6.**
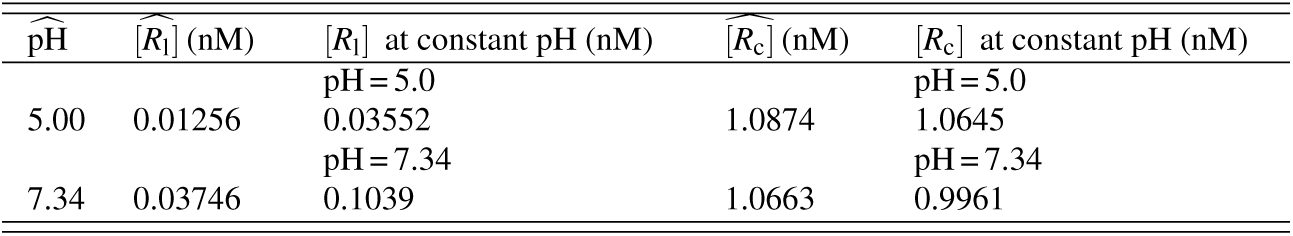
Comparison of oscillatory versus constant pH drive. The data for 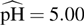 were obtained from the datafile for Fig. 3. The data for 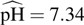 were obtained by computing a time series using the values of the parameters for Fig. 3 except 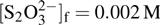.

In summary,

– The cleavage-ligation equilibrium R10 as written is driven further towards cleavage (right) under nearly-isothermal, low concentration pH cycling.

– Under high concentration, thermo-pH cycling the yield of ligated product is significantly improved because the equilibrium is driven towards ligation (left).

These results raise the possibility that, in the RNA world, nature may have made use of both regimes to tune the action of ribozymes dynamically. This example also suggests that, in the non- or proto-cellular environment of the RNA world, ribozymes which have not evolved sophisticated enzyme functionality and cannot take a reaction across a path of significantly lowered activation energy are likely to require some degree of thermal cycling, as well as pH cycling, to achieve adequate performance.

**Fig. 4.**
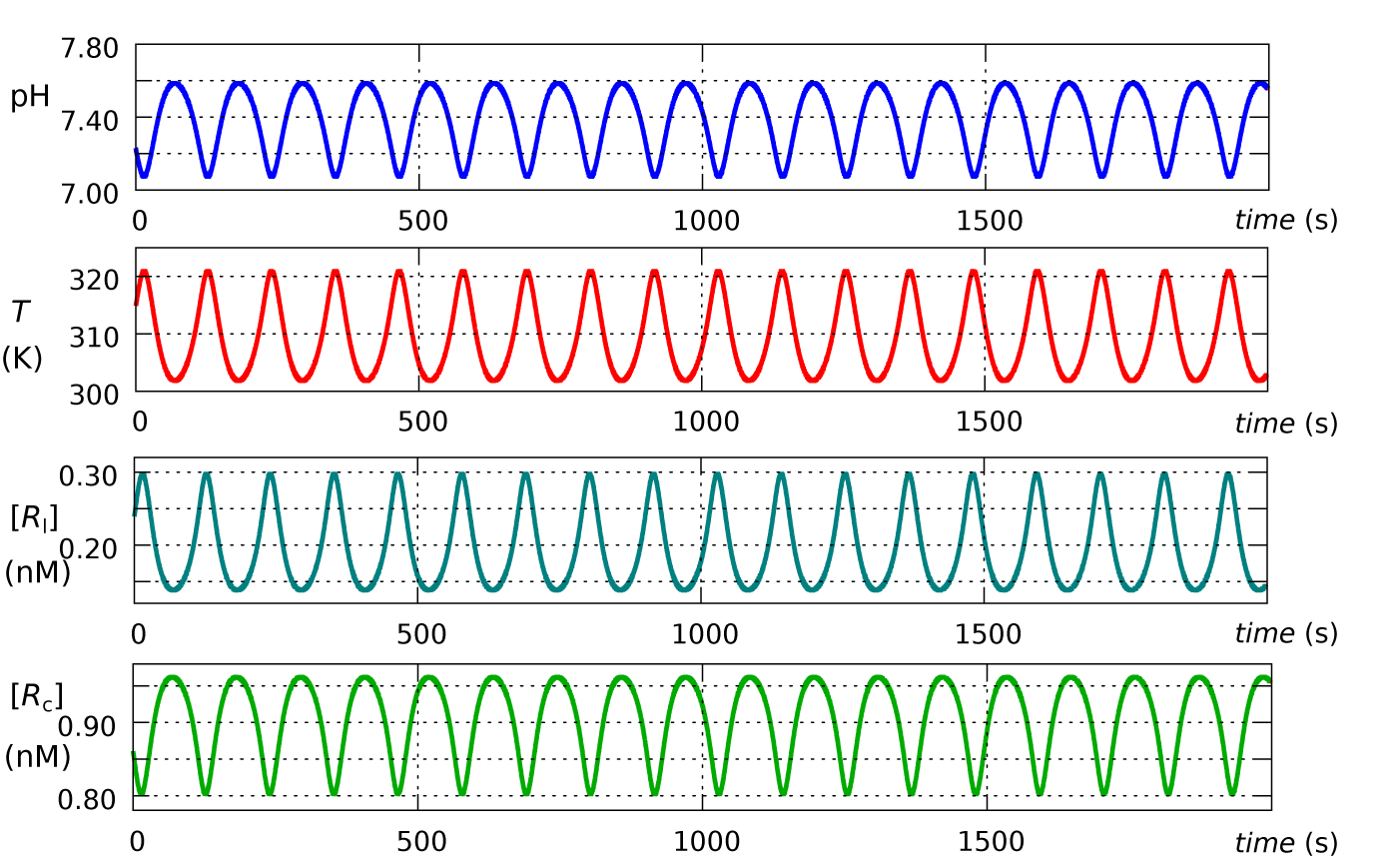
Under a mild, thermo-pH oscillatory drive, *F* = 0.44 *μ*l s^-1^, [*R*]_c,f_ = 1.1 nM, [H_2_O_2_]_f_ = 1.35 M, 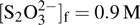, [H^+^]_f_ = 0.5 mM, 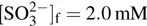, *L* = 0.002, period= 112.7 s.

**Table 7.**
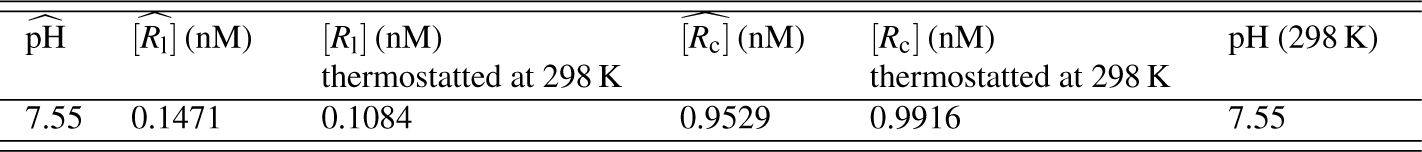
Comparison of performance under thermal and pH cycling and performance where the system is thermostatted at 298 K and the constsnt ph is adjusted to 7.55 by using [H^+^]_f_ = 0.647 mM.

## Further discussions and conclusion

From thermal oscillations to proton gradients:

The universal dependence of all life on trans-membrane proton gradients is proposed to have begun in the pH gradients across hydrothermal vents (Lane et al 2010; Lane and Martin 2012). However, this is a very narrow region. The THP oscillator can liberate the RNA world from this restrictive habitat. In a bounded, three-dimensional geometry the THP oscillator becomes a reaction-diffusion system and the oscillations manifest as travelling waves, and the RNA world could have harvested gradient energy, as life has done ever since. Quite simply, life uses proton gradients because they were there right at the beginning.

### Why not have it both ways?

There is a tension between the dual roles of RNA in the RNA world, which may be described as the ‘replication versus ribozyme activity paradox’ (Ivica et al 2013). Well-folded, stable structures are necessary for ribozyme activity, but efficient replication requires sequences that lack structure. A tradeoff between folding and templating ability means accepting only sequences with mediocre ribozyme and inefficient replication capabilities, which would more likely extinguish the RNA world than make it fitter. If the tradeoff is unacceptable, what is to be done?

Ivica et al (2013) found a structural resolution of the dilemma, where one strand of RNA could fold into a ribozyme and its complementary strand could function as an efficient replicant, through mispairing of a G to U (rather than C), which leads to the reverse-complementary, less stable AC mispair. Our results suggest that the dilemma could be resolved dynamically, a solution which avoids the necessity for specific structural conditions to propagate systematically through the RNA world, but also may cooperate with AC mispairing. In the dynamical hydrogen peroxide medium the RNA population is tightly coupled to the thermal and pH phases of the THP oscillator. A high temperature, low pH phase of the cycle may favour destructuring of a ribozyme to allow replication of some strands to take place, then replicated duplexes may denature during the next high temperature phase and adopt active ribozyme structures during a cooler, higher pH phase.

Although a dynamical resolution of the replication versus ribozyme activity paradox is a conjecture at this stage, it leads us to contemplate many more possible routes to development of the RNA world that may be enabled by thermal and pH cycling. Coupled to a dynamic thermal and pH environment, the RNA world *can* have it both ways!

### Multiple roles and versatility

One can envisage other roles for the THP oscillator in mediating proto-biotic processes, which may operate in parallel — and even synergistically — with its interactions with RNA. We have reviewed its physical properties and context in this context in Ball and Brindley (2015b).

For another example, we refer to Mayer et al (2015), who have described and tested by experiment a mechanism for vesicle formation in tectonic fault zones that relies on periodic variations in pressure to cycle carbon dioxide between the supercritical state, laden with amphiphiles, and the gaseous state, where the amphiphiles coalesce around water droplets, which settle out as bilayer vesicles into a water phase. The periodic pressure drive is postulated to be caused by tidal changes or geyser activity. A periodic temperature drive provided by the THP oscillator in the water phase could achieve much the same thing, more reliably. In this case, of course, carbon dioxide must cycle between the supercritical state and the liquid, rather than gas, state. Mayer et al (2015) postulated that the amphiphile-laden supercritical state occurs at ∼ 100 bar and 320 K. As shown in Ball and Brindley (2015a) the THP system certainly is capable of cycling between 320 K and 304 K, the critical temperature of carbon dioxide.

Still on vesicles, Lagzi et al (2010) reported on a system in which a pH oscillator controls the assembly of and rhythmically interconverts oleic acid vesicles and micelles. They used the methylene glycol-sulfite-gluconolactone pH oscillator with a period of about 2 min. In the RNA world, the THP pH oscillator may have mediated the formation of vesicles from simple amphiphiles. If so, it would also have mediated their destruction, but in a diffusive system the vesicles — along with their packaged contents — could form and move away faster than they are destroyed.

We have addressed in some depth the question of the destructive capabilities of hydrogen peroxide to nucleic acids themselves (Ball and Brindley 2014, 2015a). Here we simply note that, by the very act of oxidizing some fraction of nucleobases, hydrogen peroxide becomes an agent for change, a vector for adaptation, evolution, and diversification of RNA and its functions. In order to become a truly living system, the RNA world must replicate with fidelity — but not *too* much fidelity, and the hydrogen peroxide medium may have ensured this.

### In conclusion

We have investigated the operation of the THP oscillator on the activities of two hammerhead ribozymes, using experimental rate and thermochemical data in a coupled dynamical model for enthalpy and mass conservation. The results suggest that improved ribozyme performance can occur under oscillatory drive. For the RNA world as a whole, the dynamical environment of the THP oscillator is likely to make it fitter, resolve the dilemma of dual roles, and may help settle it into vesicles.

Finally, in efforts to detect life on other worlds a variety of characteristics of habitability, such as liquid water and an atmosphere dense enough to sustain a temperature-stabilizing greenhouse effect, and gas bioindicators, such as methane and nitrous oxide, have been proposed (Lammer et al 2009). We would add hydrogen peroxide to this list, as its presence may signal the existence of extraterrestrial RNA worlds.

**Table 8.**
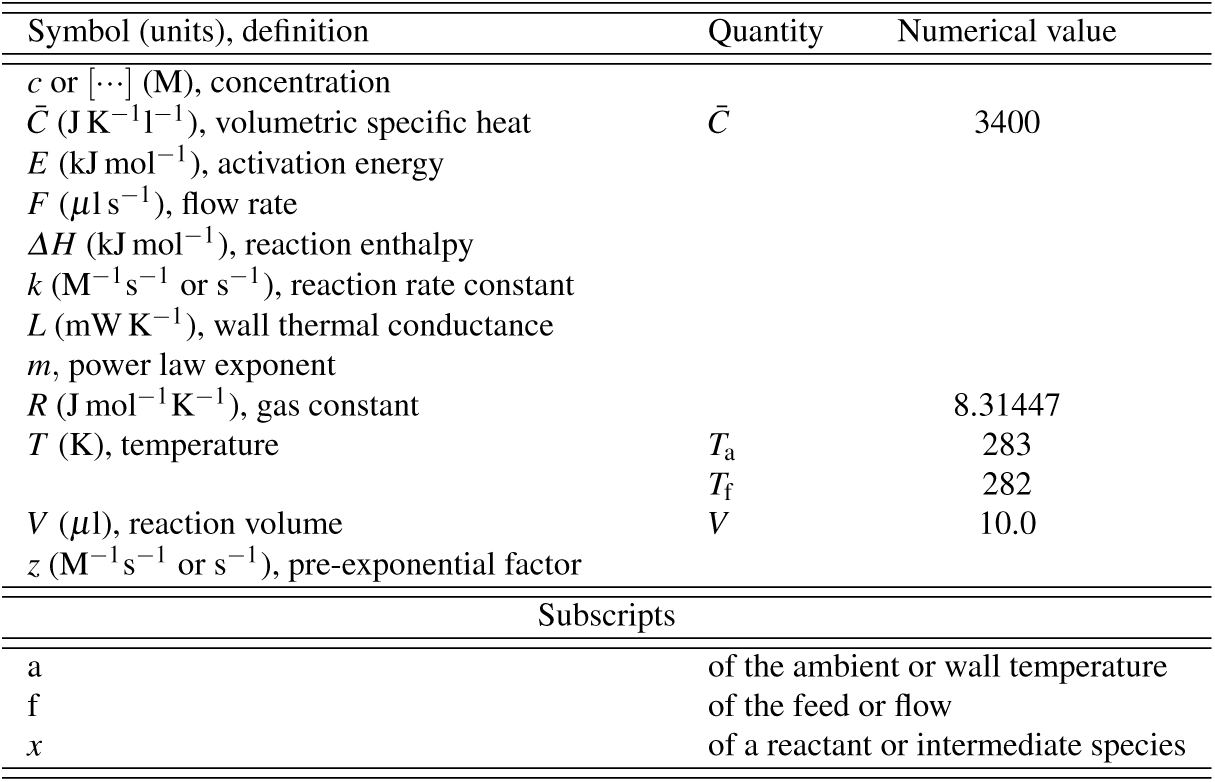
Nomenclature and numerical values.

## Acknowledgements

The authors appreciated the constructive and helpful comments and criticisms of an anonymous referee. This work was supported by Australian Research Council Future Fellowship FT0991007 (Rowena Ball).

